# Evaluating 18S Phylogenetic Placement Accuracy to Uncover Hidden Diversity in Early Branching Animals

**DOI:** 10.64898/2025.12.09.693319

**Authors:** Javier Arañó-Ansola, Inés Galán-Luque, Marc Domènech, Laura Rico-Martín, Edmund R.R. Moody, Megha Suresh, Daniel Vaulot, Javier del Campo, Mattia Giacomelli, Jesus Lozano-Fernandez

## Abstract

Small subunit ribosomal RNA (SSU rRNA) - 18S in eukaryotes - is a gene universally present across the Tree of Life and it was central in resolving ancient relationships in early molecular phylogenies. Nowadays, despite multi-locus phylogenomic approaches dominate, SSU rRNA sequences still serv as a molecular identifier of biodiversity. Phylogenetic placement uses a reference phylogeny and a given evolutionary model to annotate taxonomically environmental DNA. As such, it enables more accurate identification of divergent sequences that may belong to unknown lineages across the Tree of Life. Here we first created the largest dataset of 18S sequences belonging to non-Bilateria metazoans, by curating the sequences found in PR2 database. We then used it as a case study to test the performance of different 18S-based barcodes (V4, V9, and full-length 18S) within a phylogenetic placement framework. Employing a series of sensitivity analyses, we found that the V9 region generally lacks sufficient phylogenetic signal for reliable placements in most of the cases. The V4 region is accurate when the environmental diversity is well represented in the reference tree but struggles with divergent lineages. Full-length 18S overcome short-read limitations and emerges as the most robust option to uncover unknown major clades. Finally, we apply phylogenetic placement to empirical environmental 18S data. We observe geographical variation in non-bilaterians communities and recover a putative clade of early-branching Ctenophora based on long-sequence barcodes.

## Introduction

All cellular organisms share a structural RNA gene, the small subunit ribosomal RNA (SSU rRNA), known as the 18S rRNA gene (18S) in eukaryotes or the 16S rRNA gene (16S) in prokaryotes (Figure 1). Due to its vital role in translation, 16S/18S rRNA gene is among the slowest evolving and most conserved locus across living organisms. Thanks to this, solid hypotheses of homology for each of its nucleotides can be established among distant organisms across the entire Tree of Life (ToL) (Hillis & Dixon, 1991). Hence, it has been very useful for determining ancient evolutionary splits dating back to the Precambrian (Machida & Knowlton, 2012). As two paradigmatic examples, phylogenetic analysis using these sequences led to the influential hypothesis of the three domains of life - Eukarya, Eubacteria, and Archaebacteria - (Woese & Fox, 1977) and ushered the era of molecular systematics in zoology (Field et al., 1988).

**Figure 1:**
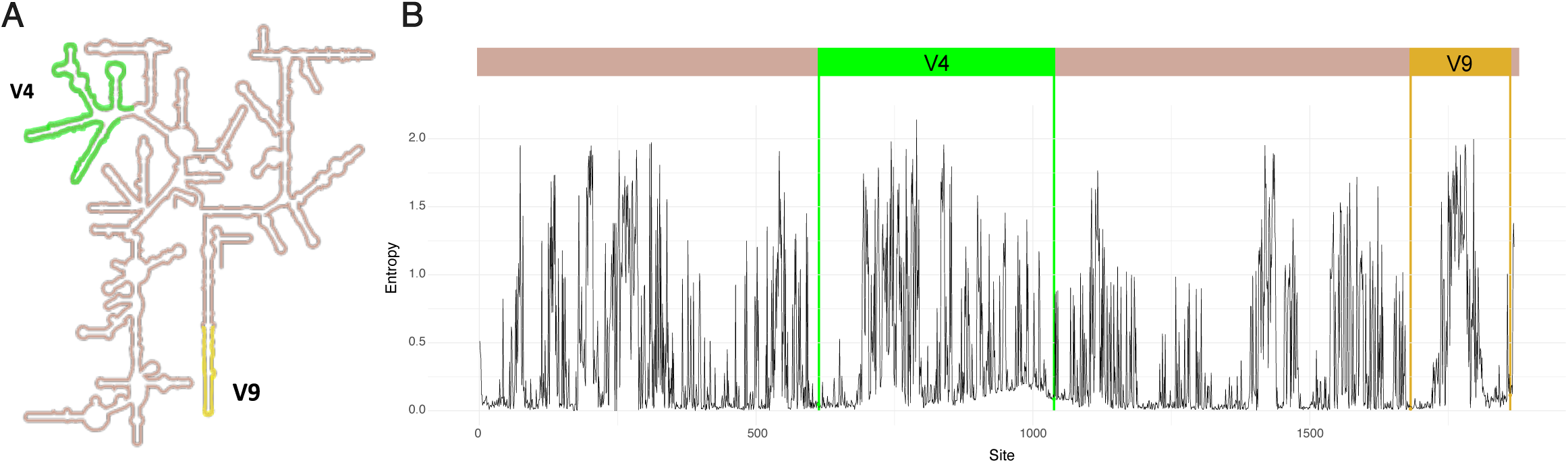
a) Secondary structure of the human 18S rRNA, adapted from RNAcentral (https://rnacentral.org/). b) Line plot showing entropy per site (i.e. the number of polymorphisms per position) in our 18S reference alignment for non-bilaterian animals. Highly variable regions V9 (ocre) and V4 (green) are highlighted, both being flanked by conserved regions in which primers bind.

As it is currently economically and technically feasible to obtain sequence data from thousands of loci (Lewin et al., 2018), the analysis of single-marker phylogenies might have become obsolete, and with them the central role of SSU rRNA in molecular phylogenetics. However, its utility is not limited to phylogenetic inference: in particular, 16S/18S rRNA barcodes are widely employed in environmental DNA (eDNA) or RNA sequencing studies. This stems from the fact that nearly universal primers can be constructed for this gene, and it is found in multiple, low-polymorphic, copies in eukaryotic genomes (Hillis & Dixon, 1991; Deiner et al., 2017; Creer et al., 2016), which usually makes it more abundant than other markers in environmental DNA. Considering these properties and its historical relevance, SSU rRNA is by far the most represented sequence in reference databases (Pierella Karlusich et al., 2023), making it the best marker for discovering previously unknown lineages across the ToL. eDNA metabarcoding is a modern non-invasive methodology for assessing community composition and structure (Bernatchez et al., 2024). The implementation of high-throughput sequencing into metabarcoding enabled the generation of large amounts of data. Nevertheless, the short-read output of most of these technologies prevents the sequencing of full-length SSU rRNAs. Instead, shorter fragments are typically amplified targeting hypervariable regions such as V4 (∼400 nucleotides) or V9 (∼150 nucleotides) (Figure 1) (Dunthorn et al., 2014). These amplicons are characterized by flanking conserved regions in which primers bind and encompass a series of phylogenetically informative sites. Recently, a new wave of long-read, high-throughput sequencing technologies has emerged (Amarasinghe et al., 2020). Their application in metabarcoding aims to overcome the long-standing trade-off between sequencing depth and the scarcity of phylogenetic signal in traditional short-read methodologies (Jamy et al., 2020).

Regardless of whether these metabarcoding studies use short or long reads, they generate thousands of genetic sequences that need to be associated with biological entities to yield meaningful ecological or evolutionary insights. Taxonomic assignment of eDNA has been traditionally based on computing pairwise similarity distances against specialized reference sequence databases (as seen in Arroyo et al., 2020; de Vargas et al., 2015; López-Escardó et al., 2018). While effective in many cases, these methods depend on arbitrary similarity thresholds that are difficult to calibrate. As a result, highly divergent sequences are often left unclassified or discarded (Santoferrara et al., 2020), leading to the loss of potentially novel diversity. Other strategies for taxonomic classification have been developed over the years (Hleap et al., 2021), including methodologies based on sequence composition (Wang et al., 2007), multinomial regression (Somervuo et al., 2016) and phylogenetics (Czech et al., 2022). Phylogenetic placement tools allocate sequences derived from metabarcoding studies (query sequences) over a predetermined phylogeny composed of reference sequences (the backbone tree), under a given evolutionary model (Czech & Stamatakis, 2019). This methodology allows taxonomic annotation of even distantly related sequences (e.g., novel groups) to reference sequences, providing a robust way to identify unknown or uncharacterised lineages from metabarcoding data. According to Czech et al. (2022), when query sequences are placed on internal branches of the backbone tree (hereafter BT), this may indicate that the sequences are difficult to classify. This can arise because the reference phylogeny lacks sufficient taxonomic representation, or because the query sequence may belong to a previously undescribed lineage or unrecognized diversity. Therefore, if the BT adequately represents all known diversity within a studied group, placement of query sequences toward inner branches indicates unknown diversity.

Here, we assembled the largest curated dataset of 18S sequences for non-bilaterian animals with two aims. First, to construct a comprehensive BT for this group. Having a trustworthy BT enriched in non-Bilateria might help identify novel or undescribed taxa, thereby reducing phylogenetic artifacts affecting early-branching Metazoa. Increasing the taxonomic diversity of phylogenetic matrices could help to resolve the root of the animal ToL, which is currently under debate (Steenwick et al., 2023). Second, to test how well different metabarcodes based on SSU rRNA work for biodiversity discovery in a phylogenetic placement framework. It remains unclear the amount of phylogenetic information that can be extracted when using the most common amplicons of the variable regions of 18S. Employing sensitivity analysis, we tested how the use of the V9 region, the V4 region, and the full-length 18S influence the accuracy of phylogenetic placement to identify known and unknown diversity. Furthermore, we tested our experimental approach to empirical metabarcoding data, including worldwide V4 and V9 samples from the Tara Oceans circumnavigation (Sunagawa et al. 2020), as well as a long-read dataset assembled by Jamy et al. in 2022.

## Materials and methods

### Reference alignment and phylogeny

While targeted SSU rRNA databases have improved significantly in recent years, they still contain errors and struggle to keep pace with the rapidly changing eukaryotic taxonomy and the influx of novel diversity (del Campo et al., 2018). Misannotated sequences in public databases hinder the assembly of high-quality reference trees needed for the accurate characterization of lineage diversity in environmental samples using phylogenetic placement techniques. To ensure an accurate non-Bilateria reference BT phylogeny, we first curated the 5,568 18S sequences annotated as non-Bilateria animals (within the phyla Ctenophora, Cnidaria, Placozoa, and Porifera) in the Protist Ribosomal Reference (PR2) database version 5.0.0 (Guillou et al., 2012). We followed the database curation pipeline described by the EukRef community. First, a set of sequences from the database was aligned to a phylogenetically accurate reference alignment. Then, a phylogenetic tree was inferred and used to identify ambiguous long branches, which may be potential artifacts, or sequences falling outside their expected clade in PR2. Following the removal of these problematic sequences, a new alignment and tree were constructed with the remaining sequences (del Campo et al., 2018). This process was iteratively repeated until no more misannotations were found. After applying the curation pipeline, we removed identical duplicates with Seqkit v2.4.0 (Shen et al., 2016). The remaining sequences were used to build the reference alignment and BT (complete BT). A set of 20 non-animal outgroup clades, including representatives of diverse eukaryotic groups, and a small representation of 32 bilaterian taxa were also included in the BT (see Supplementary Table 1).

During the curation process and the BT inference, all 18S multiple sequence alignments were inferred with ssu-align v0.1.1, which takes into account SSU rRNA secondary structure information and has been shown to achieve more accurate alignments (Nawrocki, 2009). From the alignment, phylogenetically informative sites were retained, and the uninformative ones trimmed using ClipKIT (Steenwick et al., 2020) with its dynamic threshold option. Maximum-likelihood (ML) phylogenetic trees were then inferred with IQ-TREE v2.1.2 (Minh et al., 2020), applying the GTR+G4+I substitution model. Ultrafast bootstrap was used as a measure of nodal support (Minh et al., 2013). Bayesian analyses were also performed, running four independent MCMC chains in Phylobayes-MPI v.1.9 (Rodrigue & Lartillot, 2014) using the CAT+GTR+G model, until convergence.

Reference sequences in PR2 are sometimes incomplete, lacking the hypervariable V4 and V9 regions. To avoid unnecessary computation time during the phylogenetic placement of hypervariable regions, we constructed two additional BTs, each restricted to sequences containing either V4 (V4 BT) or V9 (V9 BT). Furthermore, to ensure a fair comparison of the tested barcodes, we assembled one last BT including sequences with both hypervariable regions (common BT) that can be used to map all tested barcodes. These subsets were identified by finding in their sequence the following universal primers (Choi & Park, 2020): 5’-CCCTGCCHTTTGTACACAC-3’ (forward), 5’-CCTTCYGCAGGTTCACCTAC-3’ (reverse) for V9, and 5’-CCAGCAGCCGCGGTAATTCC-3’ (forward) and 5’-ACTTTCGTTCTTGATTAA-3’ (reverse) for V4. The number of tips included for each clade in all assembled BTs is provided in Supplementary Table 1.

### Testing phylogenetic placement accuracy

We assessed the performance of phylogenetic placement in identifying non-Bilateria diversity, both known and unknown, using three strategies.

#### Phylogenetic placement accuracy when diversity is represented in the BT

To understand the relevance of taxonomic completeness in the reference tree when mapping query sequences, we randomly removed a given percentage of sequences (tips) from each class-level clade (20%, 50% and 80%) in the BT. Then, we treated those removed tips as new query sequences. We kept at least one representative taxon from each class, allowing us to test the placement accuracy of diversity inside known classes and phyla. All barcodes were placed into the BT to evaluate its performance. As a measure of placement accuracy, we quantified the percentage of correct placement (PCP), defined as the proportion of query sequences that are confidently placed inside their clade in the BT. For all three tested barcodes, PCP was calculated for all non-bilaterian phyla and classes, and for the 20%, 50%, and 80% pruned conditions using a custom python script (Supplementary File 1). We considered sequences as confidently placed if they were assigned to their correct group with a likelihood weight ratio (LWR) ≥ 0.9, which represents the probability of the query sequence to be placed in a given branch (Czech et al., 2022).

#### Phylogenetic placement accuracy when diversity is not represented in the BT

To evaluate how phylogenetic placement would perform in the presence of novel diversity falling outside known large clades, we simulated this scenario by removing entire classes/phyla from our BTs. All pruned sequences were then treated as query sequences and placed back into the BT. We designed four scenarios in which the BT lacks major clades: (i) absence of a class from a multi-class phylum (Anthozoa and Demospongiae, from Cnidaria and Porifera, respectively); (ii) absence of a sub-phylum clade from a phylum (Medusozoa in Cnidaria); (iii) absence of a single-class phylum (Ctenophora); (iv) absence of an entire multiclass phylum (Cnidaria). For these tests, we defined as “strict correct placement” the sequences that fell in the inner branch connecting their most closely related clade present in the BT. We also defined “relaxed correct placement” when sequences were placed in any inner branch of the tree leading to any class or phylum-level clade. For both definitions, we applied different LWR thresholds (0.5, 0.7, and 0.9) for a placement to be considered correct. A schematic representation of this approach using Cnidaria as an example is shown in Figure 3A.

#### Phylogenetic placement accuracy on a biodiverse dataset

To evaluate how the tested barcodes perform when placing highly divergent query sequences belonging to groups not represented in our BTs enriched for non-Bilateria, we assembled a mock dataset by downloading 10,000 full-length reference 18S sequences with known taxonomy from the PR2 database. The rationale was to create a dataset in which the taxonomic composition was known to later compare it to the inferred taxonomy based on phylogenetic placement of V4, V9 and full 18S barcodes. This analysis aims to understand the extent to which phylogenetic placement can discriminate non-targeted sequences (non-Animals and Bilateria) from our objective taxa (non-Bilateria). We fished all V9 and V4 regions that matched the previously presented universal primers exactly, creating subsamples of the original mock dataset only containing V4 and V9 sequences with known identity. For comparability, full-length versions of the V9 and V4 mock datasets were used to evaluate the performance of full-length barcodes. Each resulting dataset was then phylogenetically placed into our BTs.

### Placing empirical metabarcoding data

We obtained metabarcode data from five different sampling sites retrieved from the Tara Ocean circumnavigation project (Sunagawa et al. 2020). Specifically, we downloaded V4 and V9 metabarcode amplifications corresponding to the size fraction of protists (project PRJEB6610 in the European Nucleotide Archive, accessed on 14 December 2024) from surface and deep chlorophyll maximum (DCM) depth layers: TARA_022 (South of Italy), TARA_058 (Madagascar), TARA_086 (Antarctica), TARA_093 (North of Chile), and TARA_151 (Azores). Long-read 18S amplicons were obtained from the dataset assembled by Jamy et al. in 2022, which contains reads encompassing the full ribosomal operon. We used Barrnap (https://github.com/tseemann/barrnap) to extract only the region belonging to 18S for each metabarcode. For preprocessing environmental reads, SolexQA++ v3.1.7.1 (Cox et al., 2010) was used with default parameters to quality-trim the reads and remove reads shorter than 100 nucleotides. PEAR (Zhang et al., 2014) was employed to merge forward and reverse reads. Exact duplicates were removed with USEARCH v12.0 (Zhou et al., 2024), and putative chimeras were identified and discarded using VSEARCH v2.28.1 (Rognes et al., 2016) with its uchime-denovo option. Finally, processed reads were clustered into operational taxonomic units (OTUs) at 97% similarity, following standard practice in environmental 18S studies. To conduct all phylogenetic placement steps, we employed Papara v2.5 (Berger & Stamatakis, 2012) to align query sequences to the reference alignment, and EPA-ng v0.3.4 (Barbera et al., 2019) to map query sequences onto the corresponding BT. OTUs placed with a LWR ≥ 0.9 on branches not belonging to non-bilaterians were automatically discarded using a custom script (Supplementary File 2). The remaining sequences were assigned taxonomic annotations based on the phylogenetic placement results using Gappa (Czech et al., 2020).

## Results

### Generation of backbone trees

From the initial set of 5,568 non-bilaterian animal sequences of PR2, 116 were flagged due to falling in incorrect positions (i.e., outside their putative clade) in the inferred ML phylogenetic trees and are currently quarantined in PR2 version 5.1.0. Additionally, 59 sequences previously annotated as class Cubozoa (Cnidaria) were found to belong to the class Staurozoa (Cnidaria), which had been absent from earlier database versions and was added in the latest release. Our curation process revealed taxonomic misannotation of 3.1% of non-bilaterian 18S sequences present in PR2. The number of sequences flagged per class, along with the reason for flagging each removed sequence, is detailed in Supplementary Figures 1 and 2. The reference alignment resulting from the curation process contained 6,567 sites, with a 22.84% completeness. After dereplication, the final BT was composed of 4,408 tips. The three additional BTs, restricted to reference sequences including either V9, V4 or both hypervariable regions, contained 883, 3,166, and 710 tips, respectively. An overview of the final BT is shown in Supplementary Figure 3 and all reference alignments and BTs can be found in Supplementary Files. In terms of topology, most well-established non-Bilaterian clades were retrieved with high support values, including Ctenophora, Cnidaria, Placozoa, as well as intra-phylum classes. Porifera, though, was non-monophyletic in the ML trees, with Silicea (Demospongiae + Hexactinellida) emerging as the sister-group to the rest of the animals. In a posterior Bayesian analysis of a taxon-reduced dataset, Porifera was recovered as monophyletic. Following the Bayesian inferred topology and current consensus in phylogenomic studies (Steenwyk & King, 2025), Porifera monophyly was forced in the common BT.

### Evaluation of 18S phylogenetic placement using sensitivity analysis

#### Phylogenetic placement accuracy when diversity is represented in the BT

When identifying query sequences belonging to known classes/phyla through phylogenetic placement, most non-bilaterian sequences were correctly annotated using the common BT (Figure 2). For the majority of the clades, PCP exceeded 75%, even when the BT lacked 80% of the group’s diversity. However, some differences existed among the tested metabarcodes. Overall, V9 performed the worst, both in terms of PCP and in the number of sequences confidently placed in the wrong group. In particular, a poor performance of this metabarcode was observed in the Homoscleromorpha sponges clade (with 100% incorrect placement) and in wrongly placing sequences into Cnidaria and Porifera. V4 performed generally better than V9, although it continued placing outgroup sequences erroneously into Porifera (Figure 2). Full-length 18S barcodes achieved a similar overall performance to V4, slightly surpassing it when querying the highly divergent Bilateria and non-animal sequences. To further understand how the amount of diversity represented in the BT affects phylogenetic placement, this experiment was repeated placing each 18S metabarcode in the taxon-richest BT available. Only the V9 region was used as a query in the V9 BT, only the V4 region in the V4 BT, and the full-length 18S sequences in the complete BT. The obtained results largely recapitulated the patterns seen using the common BT. However, the increased taxon sampling improved the correct placement results for all tested barcodes (Supplementary Figure 4).

#### Phylogenetic placement accuracy when diversity is not represented in the BT

Differences in the performance between V9, V4, and the full-length barcodes became much more pronounced when simulating the presence of unknown class/phyla among non-bilaterians (Figure 3 B). When looking at the strict correct placement, V9 analyses mostly failed to identify any of the unknown sequences, only obtaining a few correct identifications when the LWR cutoff was relaxed to 0.5. Likewise, V4 analyses showed poor results, despite performing better than V9. On the other hand, the 18S full-length resulted in better placements, albeit the percentage of correct identifications was still suboptimal for most groups (from 32% to 40% in Ctenophora, from 36.36% to 38.18% in Demospongiae, and from 40.79% to 46.0% in Anthozoa). Notably, Cnidaria and Medusozoa had an almost perfect strict placement in the common BT, even with the most stringent LWR cutoff. When considering relaxed correct placement, overall accuracy increased across all metabarcodes. Again, V9 performed worse, showing a correct placement percentage of ∼35% at best. V4, despite sensibly benefiting from the relaxed correct placement approach, only showed success rate >50% in the Cnidaria-less and Demospongiae-less scenarios when the LWR threshold was lowered to 0.5. As in the strict correct placement, full-length 18S outperformed the two hypervariable regions, with a success rate ranging from 50% in the Medusozoa-less scenario to 99.25% in the Cnidaria-less scenario under a 0.9 LWR threshold, and approaching 100% success in all scenarios with the lowest LWR threshold. According to our tests, novel classes are not inherently more difficult to identify than novel phyla among the five scenarios (Cnidaria-absent, Ctenophora-absent, Medusozoa-absent, Anthozoa-absent, and Demospongiae-absent). The experiment was, again, repeated using the taxon-richest available BT for each barcode, providing similar patterns (Supplementary Figure 5).

#### Phylogenetic placement accuracy in assessing clade richness on a mock biological dataset

When the mock dataset was analysed to infer the relative richness of each non-bilaterian class, V9 was completely unable to capture the real diversity in the sample, even when using the taxon-richest V9 BT (Figure 4 B). The shortest metabarcode provided a skewed picture of real diversity due to its extreme overrepresentation of non-bilaterian reads in the sample (49.97% and 48.31% inferred with the common BT and with the V9 BT respectively, compared to only 1.39% in the actual dataset). In contrast, the V4 region provided more accurate results, although specially influenced by the BT used. When placed in the common BT, V4 kept overrepresenting non-Bilateria (6.14% inferred vs 2% real), with most misidentification being placed among the long-branching clades Hexactinellida and Ctenophora. However, when the taxon-richest V4 BT was employed, V4 barcodes were able to capture the diversity hidden in the mock dataset with only minor errors, effectively discarding most of non-animal and Bilateria taxa (2.62% of non-Bilateria inferred vs 2% real) Remarkably, when using the taxon-richest BTs, V4 outperformed the full-length 18S metabarcode (Figure 4 B). To further assess whether the low phylogenetic signal of V9 was simply due to its reduced length, the V4 amplicons from our mock dataset were divided into three subfragments of 175 nt (to mimic the mean length of the V9 amplicons used in this experiment) and were placed as query sequences in the V4 BT. None of the V4 subfragments correctly reconstructed the taxonomy of the artificial community on their own. The inflated representation of non-bilaterian sequences observed with V9 amplicons was also detected with shortened V4 fragments, though to a lesser extent (Supplementary Figure 6).

**Figure 2:**
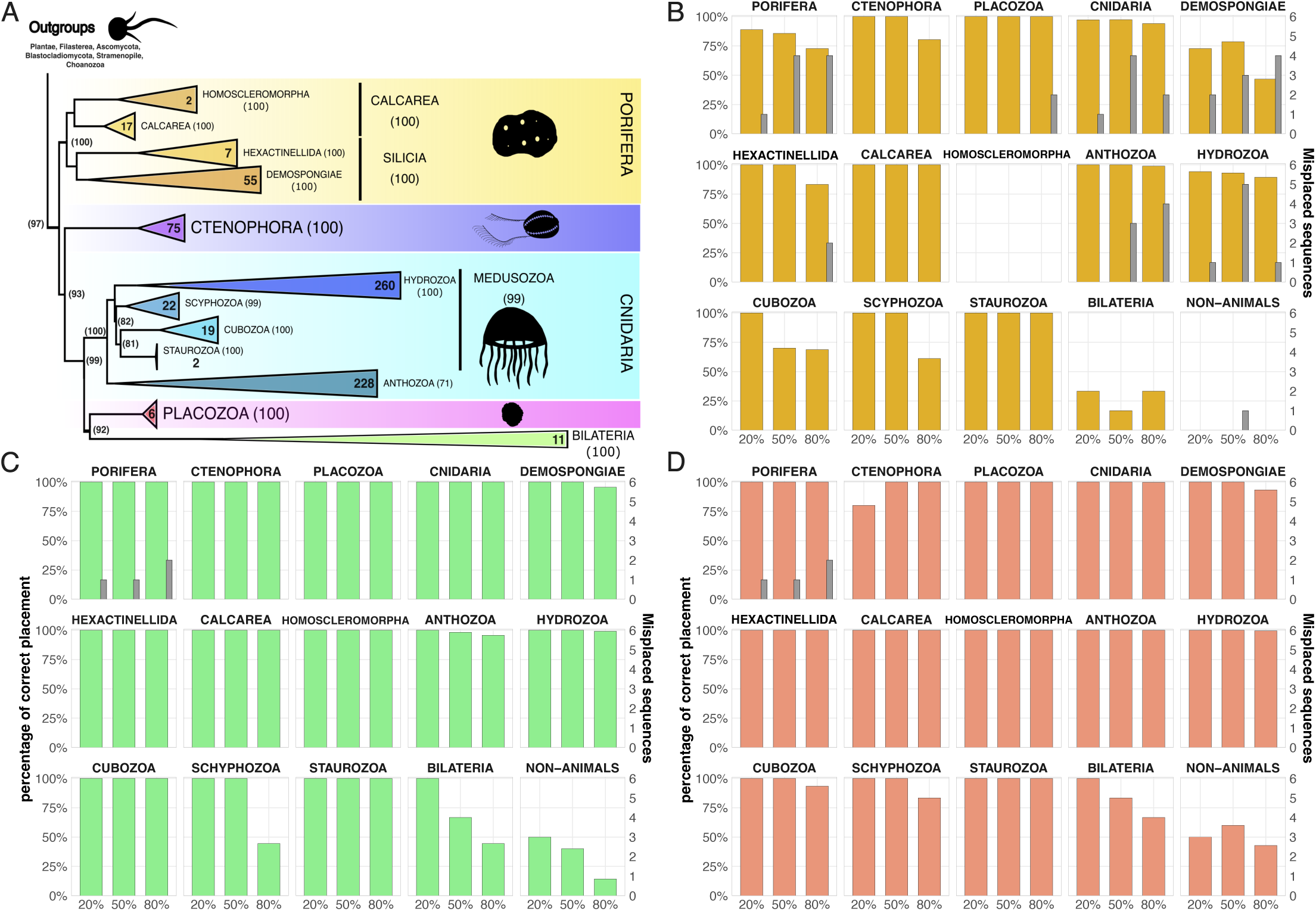
Comparison of placement reliability of 18S-based barcodes across different classes and phyla represented in the backbone tree (BT). A) Topology of the common BT used to compare the phylogenetic placement reliability of each tested barcode. Ultrafast Bootstrap support values are shown in brackets and the number of tips inside each collapsed clade is indicated in bold. Silhouettes for animal groups were obtained from the public domain https://www.phylopic.org. B), C), and D). Stacked bar charts plotting the PCP (left y axis) resulting from the phylogenetic placement of V9, V4 and full-sequence 18S barcodes, on different percentages of BT completeness (x axis). Overlaid histograms represent the number of query sequences confidently placed in the wrong clade (right y axis).

**Figure 3:**
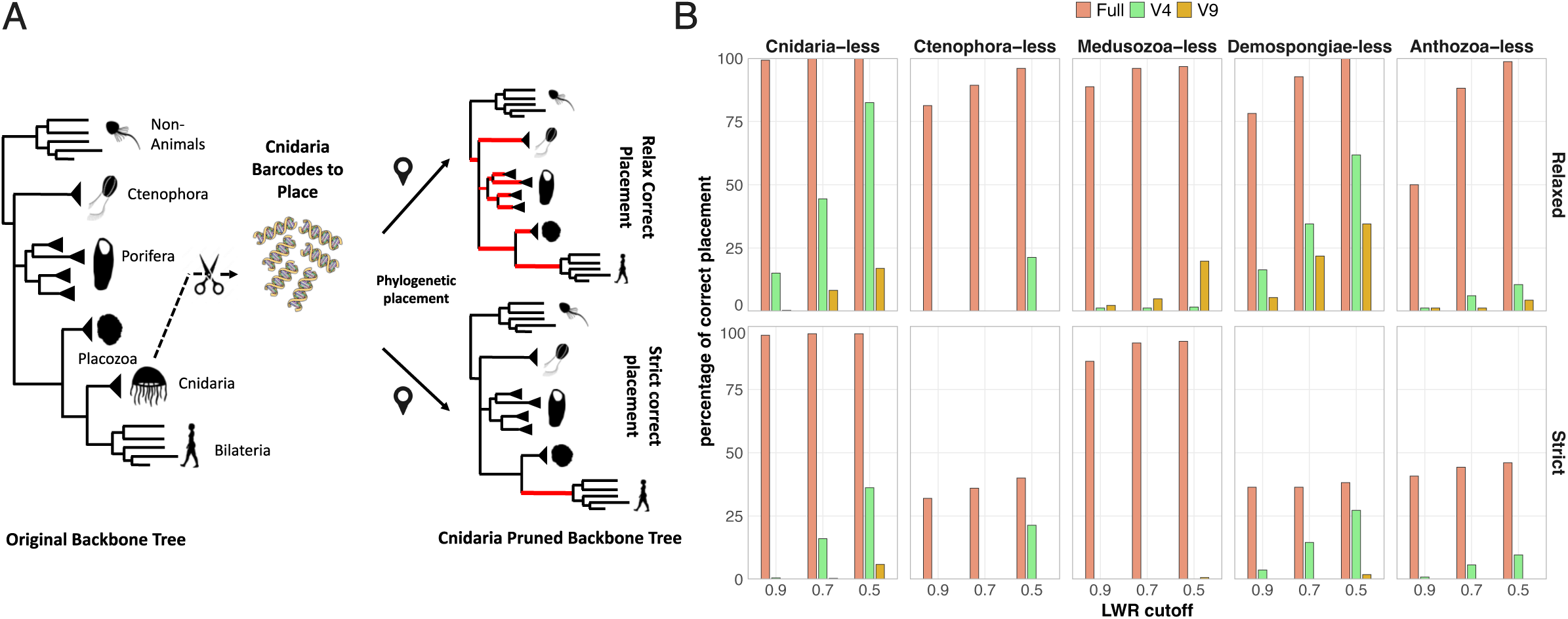
A) Graphic representation of *strict* and *relaxed* correct placement introduced to test the behaviour of phylogenetic placement when a clade is not represented in the BT. Silhouettes for animal groups were obtained from the public domain https://www.phylopic.org. B) Histogram showing PCP of 18S-based barcodes belonging to hypothetical unknown classes/phyla. Different LWR thresholds are considered for a placement to be counted as correct (x axis). Top: results under the relaxed correct placement. Bottom: results under the strict correct placement.

### Application to real metabarcoding data

V9 and V4 metabarcoding data from five Tara Oceans stations spread across different ocean basins were analysed (Figure 5) always using the taxon-richest BT available for each barcode. The discrepancies between V9 and V4 seen in the mock community experiment were also evident here: the proportion of V9 reads annotated as non-bilaterian animals was often an order of magnitude higher than their V4 counterparts. Moreover, the class-level richness inferred from V9 remained constant across all sites, mirroring the same results seen in the V9 portion of Figure 4. Given our simulation results, we focus on V4 data to unveil the distribution patterns of known non-bilaterian clades, while full-length 18S metabarcoding was used to explore the potential presence of undescribed classes or phyla near the base of the animal ToL. Non-bilaterians accounted for 0.4-2.5% of OTUs in the five Tara Oceans sites explored here, with diversity patterns appearing to be mostly driven by geographic location rather than depth. Cnidaria generally emerges as the most diverse phylum, mainly due to the class Hydrozoa, followed by Porifera and Ctenophora. However, in some instances, Porifera is as diverse as Cnidaria or even becomes the dominant clade in terms of OTU diversity, as observed in the Azores and Antarctica samples.

As long-read data was not available from the Tara Oceans stations, a recently published dataset that includes eDNA from multiple expeditions, extending down to the bathypelagic depth layer, was examined (see Supplementary Table 2 for sampling regions). In this dataset, five OTUs were confidently placed as a putative unknown lineage, branching as the sister group to all known Ctenophora sequences. All these sequences were recovered exclusively from the bathypelagic layer (1200 m depth) at the 49th sampling site of the Malaspina circumnavigation (Duarte, 2015). To further evaluate the identity of our putative deep-branching ctenophores, BLAST searches were performed against the NCBI database. This analysis aimed to discard sequences with high BLAST score similarity (>95%) to an already existing sequence not belonging to a ctenophore (excluding terms like “uncultured eukaryote” or “unknown eukaryote/Metazoa”). Two of the five putative novel ctenophores identified with phylogenetic placement had a best BLAST hit against a ctenophore (90.58% and 93.78% similarity). Two others had slightly more similar matches to non-metazoan taxa but also showed close hits with a ctenophore (90.40% ctenophore against 92.18% dinoflagellate, 89.61% ctenophore against 90.19% Rhizaria). The remaining OTU had its best hit (96.1%) to Alveolata and only 87.8% to ctenophores, and was therefore discarded. In order to corroborate the phylogenetic position of these five sequences as early branching ctenophores, a ML phylogenetic tree was built, including the candidate uncharacterised ctenophores sequences alongside a set of reference 18S sequences. These environmental OTUs formed a clade sister to all known ctenophores, with full ultrafast bootstrap support (Figure 6). A phylogenetic analysis within a Bayesian framework was also conducted. Due to convergence issues, we had to use a reduced version of the 18S dataset that we had previously used for the ML tree. The only topological discrepancy between the two methods involves the monophyly of Porifera, which is only recovered as a clade with the Bayesian approach. The position of the environmental ctenophores sequences remained stable in both methods as sister to all other sequenced 18S ctenophores.

**Figure 4:**
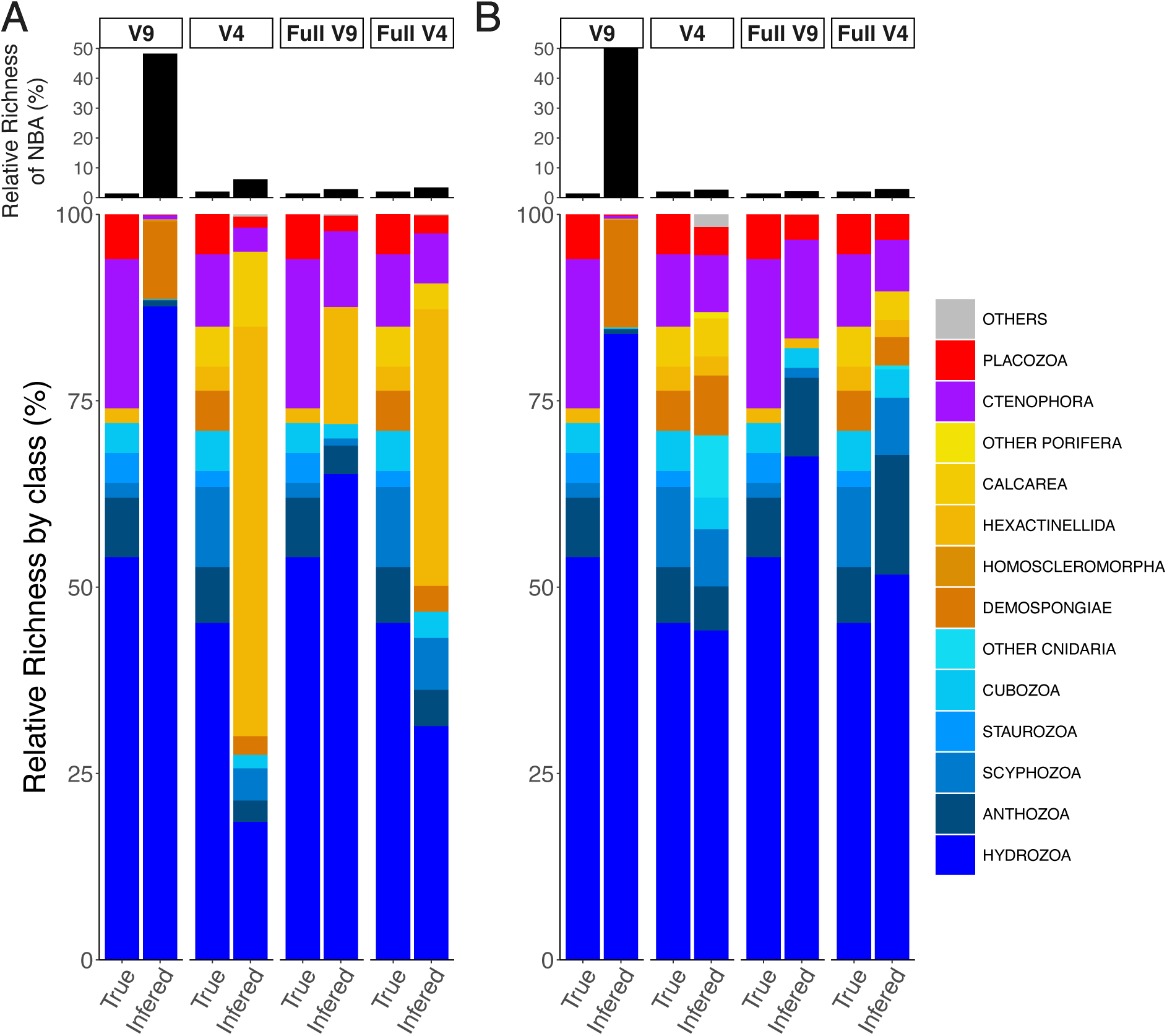
Stacked bar charts showing the taxonomic analysis of the assembled mock community using different 18S barcodes. For each barcode, the true taxonomic identity of the community is compared to the inferred identity using phylogenetic placement and the given barcode. Both the percentage of OTUs assigned as non-bilaterian animals (NBA) over the total number of OTUs (top), and the relative proportion of each non-bilaterian class/phyla (bottom) are shown. A) Inferred taxonomy of the mock dataset placing queries in the common BT. B) Results inferred when placing queries in the taxon-richest BT for each barcode. “Full V9” and “Full V4” refers to full length versions of the V9 and V4 mock datasets. This allows for a fair comparison between full-length 18S barcodes and each of its short-read counterparts.

**Figure 5:**
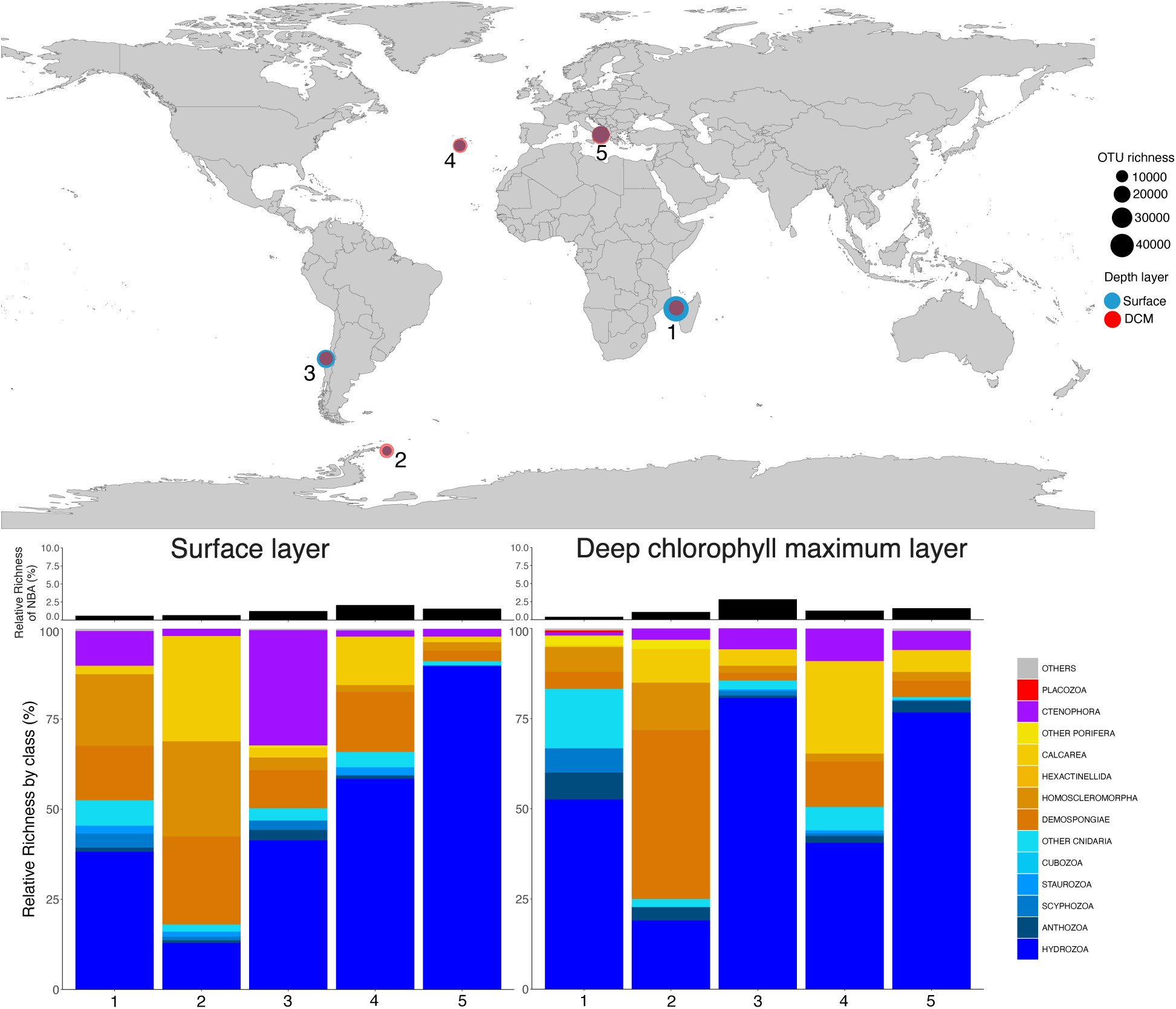
Overview of the phylogenetic placement analysis of V4 metabarcoding data from five Tara Ocean sampling sites. The map at the top displays the location of the studied Tara stations; dot size is proportional to the number of OTUs sampled at each site. At the bottom, the percentage of OTUs assigned as non-bilaterian animals (NBA) over the total number of OTUs, and the relative proportion of each non-bilaterian class/phyla are shown, both for the surface water layer (left) and the DCM layer (right).

Although this work has focused on the study of 18S, the environmental sequences from which we detected our divergent ctenophores contain information about the whole ribosomal operon. Accordingly, an additional ML phylogenetic tree based on the 28S gene was inferred. Sequences extracted from the ribosomal operon database (Krabberød et al., 2024), as well as the suspected ctenophores were included in the analysis. Contrary to the 18S results, the 28S tree placed the environmental sequences nested within crown Ctenophora, showing a discordant evolutionary story between the two ribosomal RNAs.

**Figure 6:**
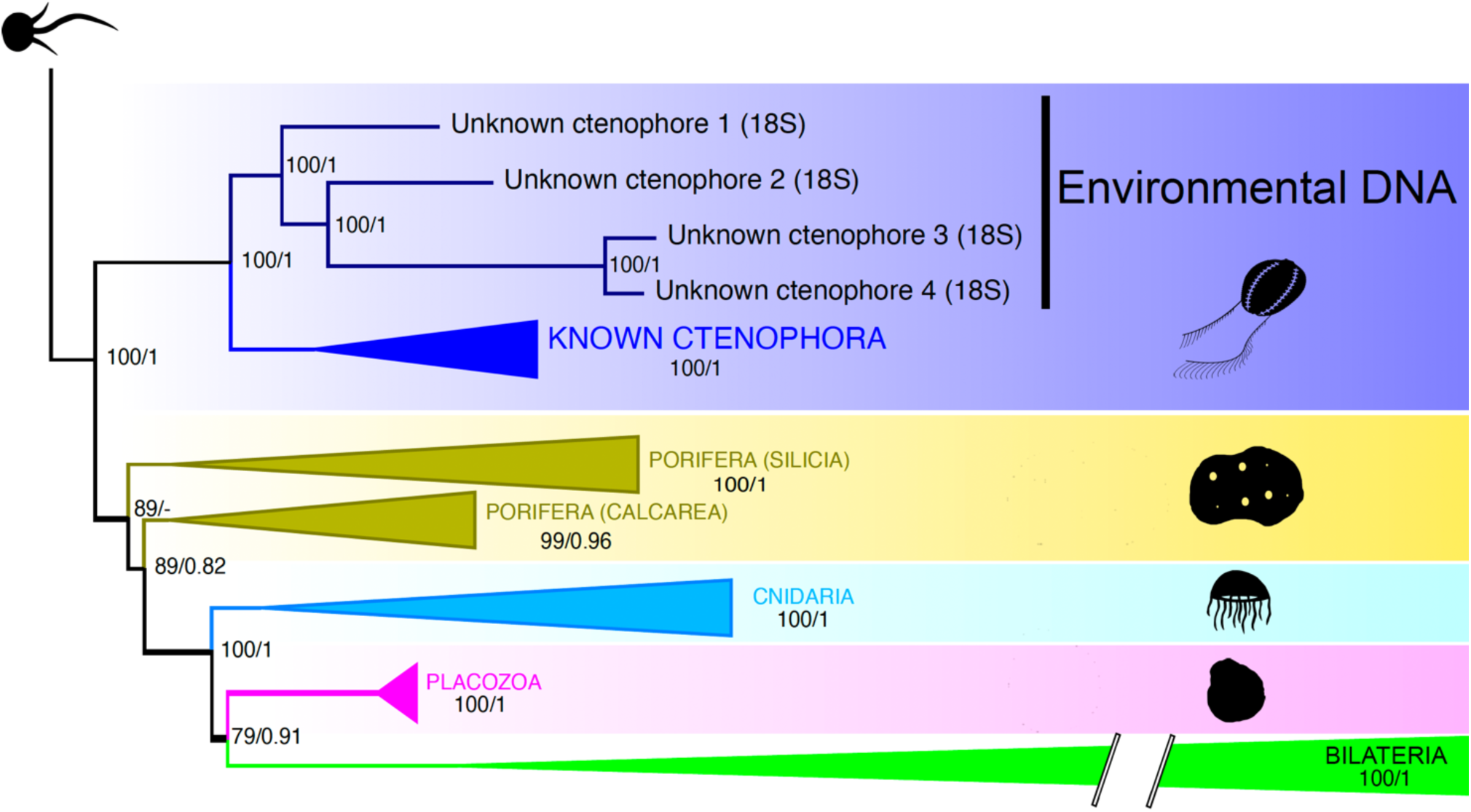
Collapsed phylogenetic tree inferred from 18S sequences spanning all animal classes, together with environmental OTUs identified as early-branching ctenophores in the phylogenetic placement analysis of Jamy et al. (2022) long-read 18S metabarcoding dataset. The topology shown corresponds to the ML analysis. Support values are reported as ultrafast bootstrap / posterior probability (from the Bayesian analysis). Nodes lacking posterior probabilities indicate a disagreement between the two methodologies. Silhouettes for animal groups were obtained from the public domain https://www.phylopic.org. The Bilateria branch is not up to scale.

## Discussion

We assembled the largest curated reference 18S dataset for non-bilaterian animals in the database PR2, comprising thousands of sequences spanning all known molecular diversity of this group. While López-Escardó et al. (2018) had already generated a phylogenetically curated metazoan 18S reference dataset, their collection was primarily focused on the more diverse bilaterian animals, with over half of the entries in the dataset constituted by arthropods. We inferred three well-supported ML phylogenetic trees, each tailored to a different type of 18S barcode. These will serve as *bona fide* BTs in which novel 18S can be mapped through phylogenetic placement. During the BT assembly, we identified several misannotations among the sequences classified as non-bilaterian animals in the PR2 database. If such errors occur at the class and phylum levels, they are likely even more common at shallower phylogenetic scales, such as order or family. This highlights the importance of deeper curation efforts in major sequence repositories across the ToL, such as those being carried out by the EukRef community within the PR2 database.

### Evaluating 18S barcodes for phylogenetic placement

Significant work has compared the usefulness of analysing amplicons from the hypervariable regions V4 and V9 across several biological groups (Choi & Park, 2020; Tragin et al., 2018; Stoeck et al., 2010; Piredda et al., 2017). Most studies report discordant results between the two markers when analysing the same community, both in terms of taxonomic annotation and the number of reads or OTUs recovered. Generally, V9 can detect rare taxa that V4 fails to reveal. Explanations for this discordance have focused on technical issues during sequence sampling, including differential sequencing error rate (Tanabe et al., 2016) and primer bias (Silverman et al., 2021), as well as biological problems related to intracellular 18S diversity (Decelle et al., 2014). However, the majority of these studies rely on similarity-based approaches for taxonomic annotation. The performance of each barcode within a phylogenetic framework has been less explored.

Our results caution against the use of the V9 region for phylogenetic placement, at least for non-bilaterian sequences. Taxonomic annotations based on V9 reads may retain some credibility when most of the reads belong to taxa well represented in the BT. Otherwise, the V9 region lacks enough phylogenetic signal to confidently place highly divergent sequences, which sometimes appear spuriously placed distantly from their true relatives. This is neatly illustrated in Figure 4, in which V9 region analyses show poor performance, as well as in Figure 2, with the low percentage of confidently placed V9 outgroup sequences. V4 amplicons outperform their V9 relatives in all our tests. Durthorn et al. (2014) already stated that including V4 amplicons in an alignment of full-length reliable 18S sequences represents the most appropriate approach for using high-throughput sequencing data to answer phylogenetic questions in ciliates. When we trimmed V4 into shorter portions equivalent in length to V9, V4 fragments yielded poor diversity estimates, although closer to reality than the V9 fragment. This suggests that the poor performance of V9 is attributable not only to its short length but also to a lack of phylogenetic informativeness of this region in our group of organisms. Full-length 18S was by far the best marker when testing phylogenetic placement of taxa not represented in the reference BT, indicating that long-read sequencing can increase the accuracy in associating divergent OTUs to characterized organisms in metabarcoding studies. The idea that the ability of a molecular phylogenetic marker to recover the correct tree increases with its length is neither new nor revolutionary (see Page & Holmes, 2009). However, understanding and determining the threshold at which a given amplicon ceases to provide a reliable signal at particular taxonomic levels is crucial for deciding whether long-read sequencing is necessary for the question at hand. For biodiversity discovery-particularly for detecting unknown lineages-, longer, full-sequence, 18S barcodes appear the most suitable marker, as traditional, shorter, NGS amplicons consistently underperform. Yet in some cases, even when a complete 18S was available, some highly divergent query sequences belonging to classes or phyla absent from the BT could not be confidently placed. This implies that hidden biodiversity, especially the one including highly divergent lineages, may often be underestimated in phylogenetic placement studies, even when using long-read 18S metabarcodes. Moreover, the same set of experiments performed both in the common BT and in barcode-specific BTs, highlights the importance of a good reference taxon sampling not only for the targeted group but also for the outgroups. Indeed, the difference in outgroup representation is the most plausible explanation for the discordance in accuracy seen in the analysis of our mock community using V4 barcodes between the common BT (six non-animal taxa, 11 Bilateria taxa) and the taxon-richer V4 BT (16 non-animal taxa, 21 Bilateria taxa). Given a low representation of non-targeted sequences in the BT, divergent queries belonging to those non-represented taxa tend to be placed towards long branching ingroup branches. Although our results are limited to phylum/class-level clades in non-Bilateria, further research exploring shallower taxonomic classification, other kinds of metabarcodes, and other biological groups may benefit from the resources provided here.

### Exploring the known and unknown diversity of non-bilaterians

The group here referred to as non-bilaterians is paraphyletic and extremely diverse, representing multiple animal phyla with distinct evolutionary paths (Giribet & Edgecombe, 2020). Differences in ecology, physiology, and morphology among these animals make it challenging to identify shared distribution and diversity patterns. Most metabarcoding studies assessing metazoan distribution patterns have treated non-bilaterian phyla as part of a broader animal dataset, without specifically focusing on these early branching animal clades (Leray & Knowlton, 2015; López-Escardó et al., 2018; Geraldi et al., 2024). Other works target particular non-bilaterian clades (Sun et al., 2025; McCartin et al., 2024). While the sample size analyzed here is too limited to draw any conclusions about the global distribution of non-bilaterians, our results provide a robust framework and a set of resources for future research. Notably, the proportions observed in our phylogenetic placement of five different sites from the Tara Oceans expedition are in some concordance with what has already been reported in other global animal metabarcoding studies. Cnidaria have often been found as one of the most predominant phyla in terms of diversity (Clarke et al., 2021; Geraldi et al., 2024; López-Escardó et al., 2018), and Verhaegen et al. (2025) recently observed, using the COI marker in Arctic regions, that Porifera exhibit elevated diversity in deep, cold waters. We clustered metabarcodes at 97% similarity, following the conventional threshold for grouping sequences into the same OTU (Stackebrand & Goebel, 1994; López-Escardó et al., 2018). We acknowledge, however, that 97% similarity clustering is not a reliable proxy for species-level classification in most eukaryote groups, and it may be underestimating diversity. Indeed, we observed that in Ctenophora, Calcarea, Placozoa, Scyphozoa, and Staurozoa, most of the sequences included in our curated dataset diverge by less than 3%, which highlights that our diversity estimates regarding these groups may be underestimating true species richness (Supplementary Figure 4 B). We nonetheless followed a common clustering practice since testing the optimal threshold level for species identification is out of the scope of this work. Phylogenetic placement of long-read metabarcoding data allowed us to identify a potential clade of ctenophores branching as the sister group to all known Ctenophora sequences. This finding is particularly exciting for understanding early animal evolution, as the addition of intermediate taxa along long branches —such as those leading to Ctenophora— can help mitigate long-branch attraction artifacts in phylogenetic inference (Lartillot et al., 2007). However, a 28S phylogenetic tree placed these environmental sequences nested among known ctenophores, remarking the need for a multi-locus or specimen-based validation. We acknowledge that the signal of the 18S sequences could be due to sequencing errors related to long-read sequencing (e.g., chimeras) or even long-branch attraction artifacts. Nonetheless, multiple well-known clades in protistology and microbiology were first identified by eDNA (van Hannen et al., 1999; Kolodziej et al., 2007; Howe et al., 2011), including the archaeal clade shown to be sister to all eukaryotes (Spang et al., 2015). Future research may integrate information from the ribosomal operon to build a robust BT able to identify sequences based on the full 18S, 28S, and ITS sequences, as whole operon metabarcodes may be the direction towards ribosomal-based metabarcoding is moving.

## Conclusions

In the rise of the genomics era, we turn our attention back to a gene that was pivotal in understanding the ToL as we do nowadays. We contribute to the curation of SSU rRNA genetic resources and test its value in animal diversity discovery through phylogenetic placement. We also provide four reference BT for non-bilateral animals. Our results, based on the analyses of animal sequences, advocate for using the V4 region in metabarcoding projects. We found this region as more reliable than the V9 for identifying classes or phyla included in the reference tree, yet showing problems when dealing with unknown clades. These limitations can be partly solved using full-length 18S barcodes. When applying phylogenetic placement to real empirical data from five different worldwide marine locations, we saw geographic variation in the analysed non-bilaterian communities, with Cnidaria or Porifera alternating dominance. Additionally, we gathered evidence of an unknown or uncharacterised early-splitting clade of likely deep-sea ctenophores based on full-length 18S metabarcoding data. By clarifying the potential and limitations of 18S amplicons in phylogenetic placement, this study provides a robust framework for using environmental sequencing to uncover hidden biodiversity across the ToL.

## ACKNOWLEDGEMENTS

JL-F was funded by Ministerio de Ciencia e Innovación of Spain and Next Generation EU (MCIN/AEI/10.13039/501100011033; grants ‘Ayudas para Incentivar la Consolidación Investigadora’ CNS2022-135805 and PID2022-137753NA-I00), and Comissió Interdepartamental de Recerca i Innovació Tecnològica (2021SGR00279). Authors JA-A, IG-L, and LR-M were also supported by FI and FI-STEP contracts from the Catalan Government (2025 FI-1 00443, 2024 FI-1 00804, and 2025 STEP 00397, respectively). The author ERRM was endorsed by a Marie Skłodowska Curie Postdoctoral Fellowship with code 101147105 under HORIZON-MSCA-2023-PF-01.

## Supplementary

**Supplementary Figure 1:**
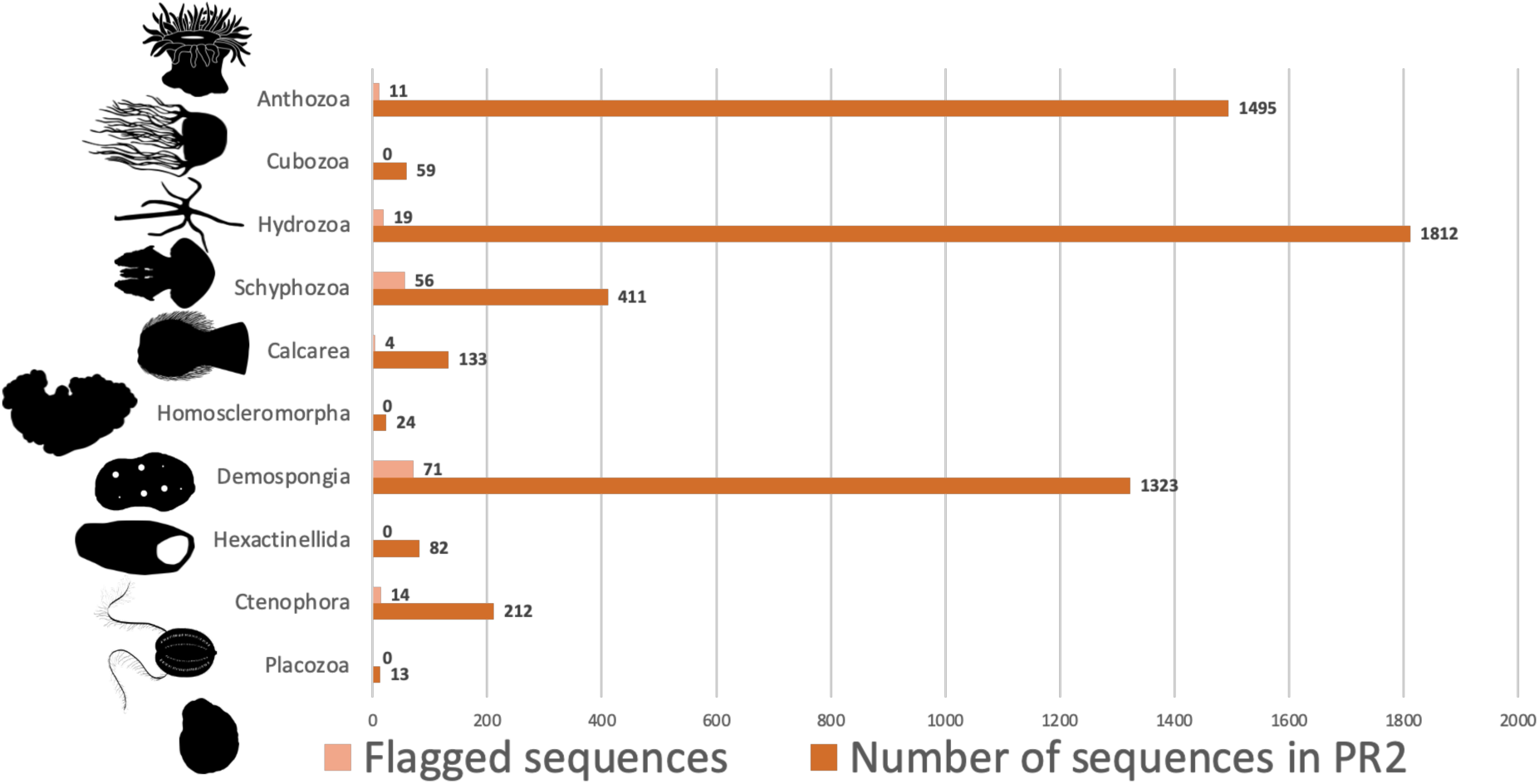
Number of current sequences of each non-bilaterian Metazoa class in the PR2 database version 5.0.0 compared to the number of sequences flagged during the curation process.

**Supplementary Figure 2:**
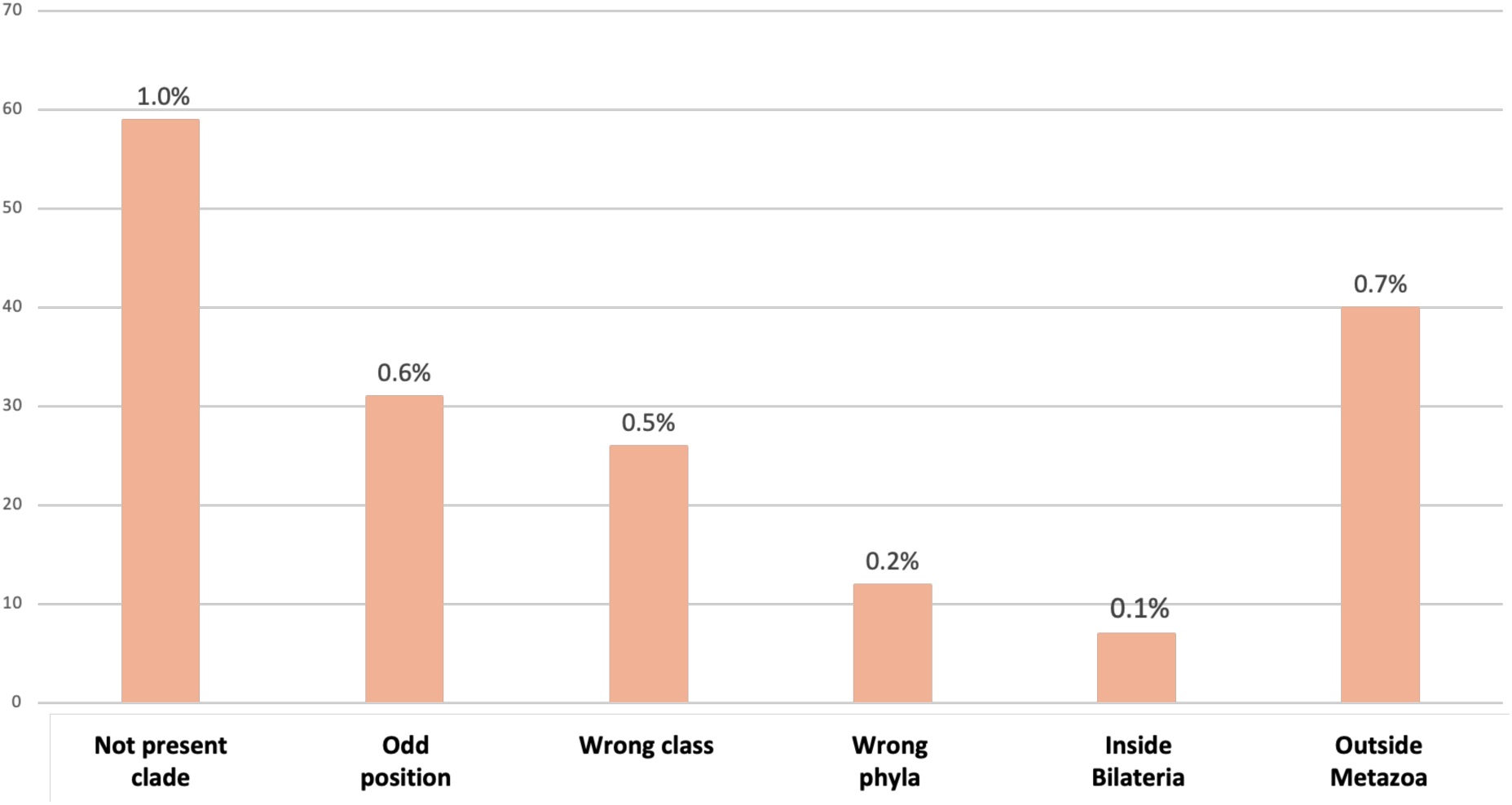
Reason for flagging each problematic sequence during the curation process of non-bilaterian animal taxa in the PR2 database (percentage over total non-bilaterian animal sequences in PR2).

**Supplementary Figure 3:**
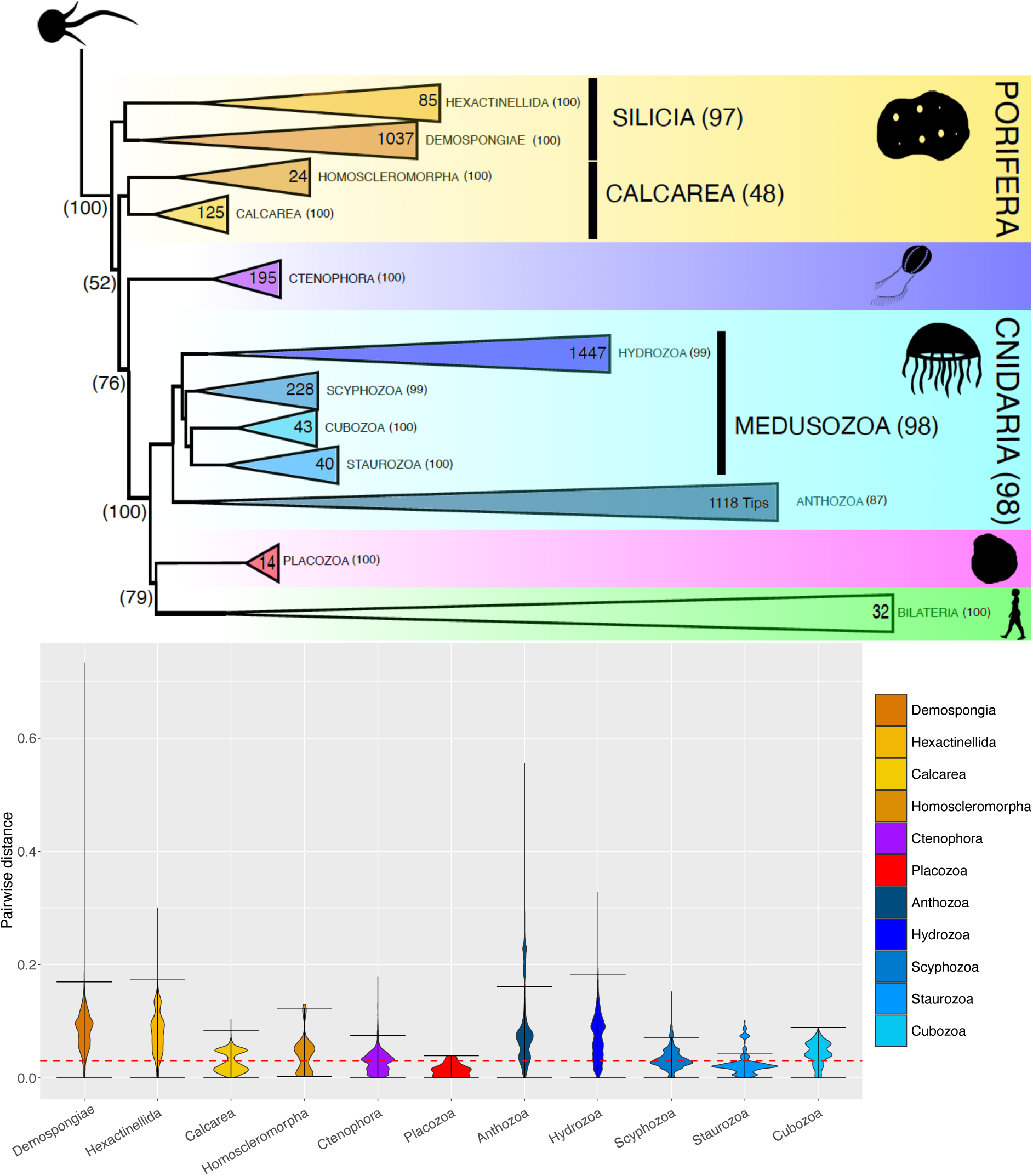
A) Collapsed ML backbone phylogenetic tree containing all the 18S sequences in the PR2 database belonging to non-bilaterian animals after erasing all problematic sequences (bipartition support assessed with ultrafast bootstrap). The number of sequences of each clade can be seen inside each collapsed node. B) Violin plot depicting pairwise distance between all 18S sequences within each class from the complete backbone tree. Pairwise distances were computed from our reference alignment. Any position with at least one of the compared sequences presenting a gap was not included in the calculation. The dashed red line highlights the 0.03 pairwise distance threshold below which sequences would be clustered under a 97% similarity clustering algorithm.

**Supplementary Figure 4:**
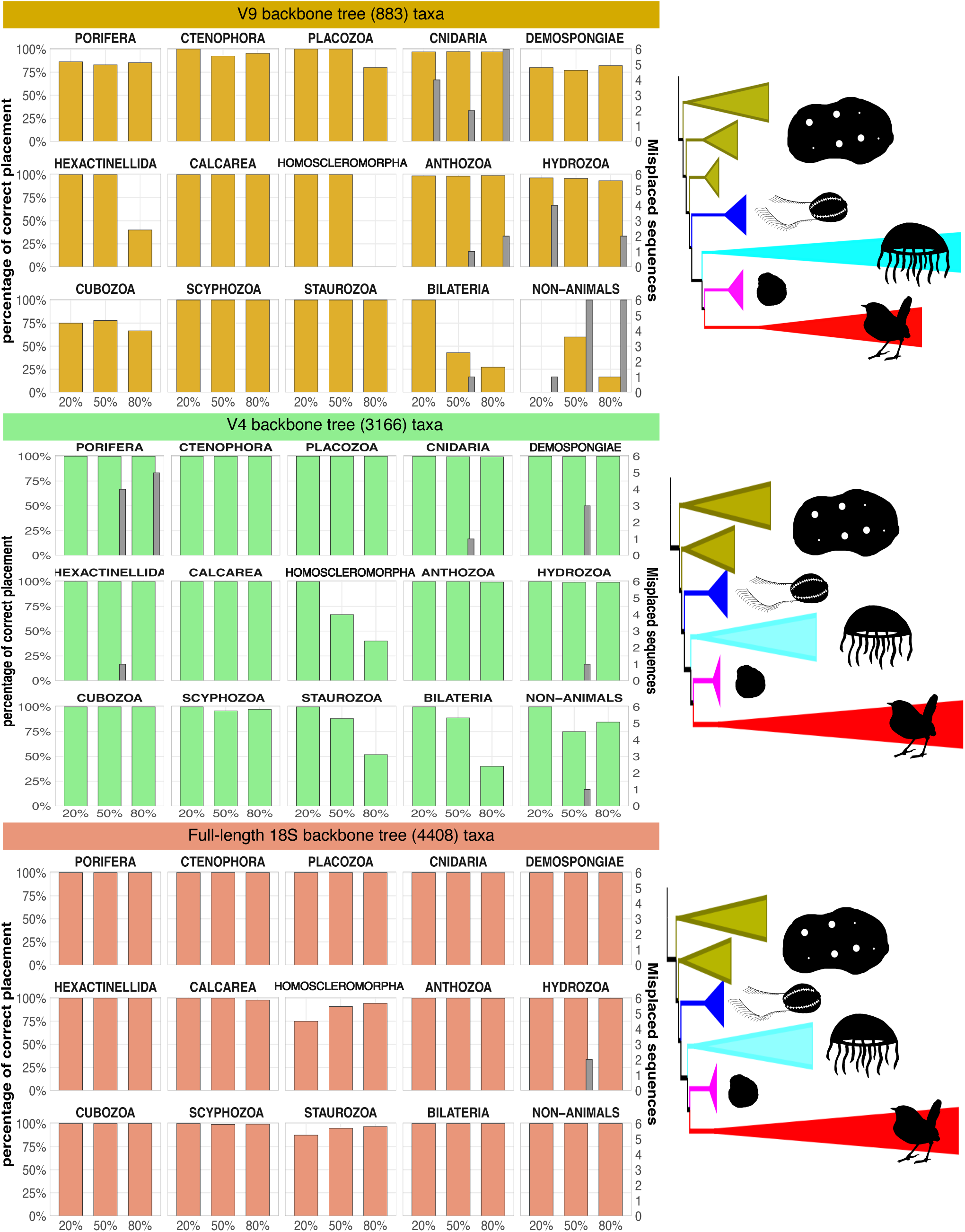
Comparing placement reliability of 18S-based barcodes belonging to different class/phyla represented in the BT, using the taxon-richest BT available for each tested barcode. The stacked bar chart plots PCP (left y scale) for each of the tested barcodes and levels of completeness of the BT (x axis). The intercalated histograms represent the number of query sequences (right y axis) confidently placed in the wrong clade. An overview of the topology of each BT, displaying the main non-bilaterian clades, is shown on the right side.

**Supplementary Figure 5:**
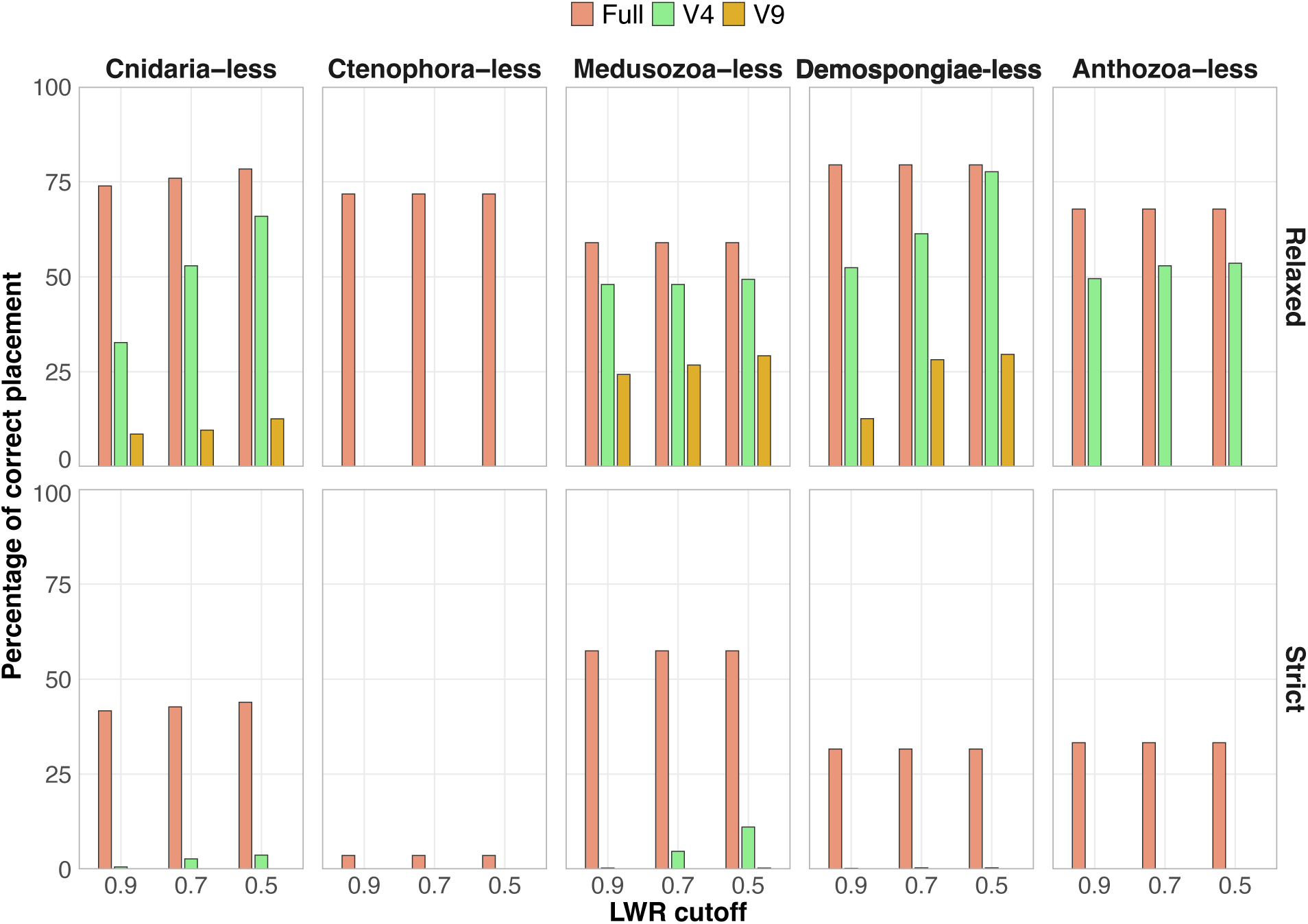
Histogram showing PCP of 18S-based barcodes belonging to hypothetical unknown classes/phyla. Different LWR thresholds are considered for a placement to be considered correct (LWR cutoff). Also, different placement precision relative to the branch of the BT that represents correct placement are recognized (Relaxed and Strict correct placement). The taxon richest BT available was used for the phylogenetic placement of each barcode.

**Supplementary Figure 6:**
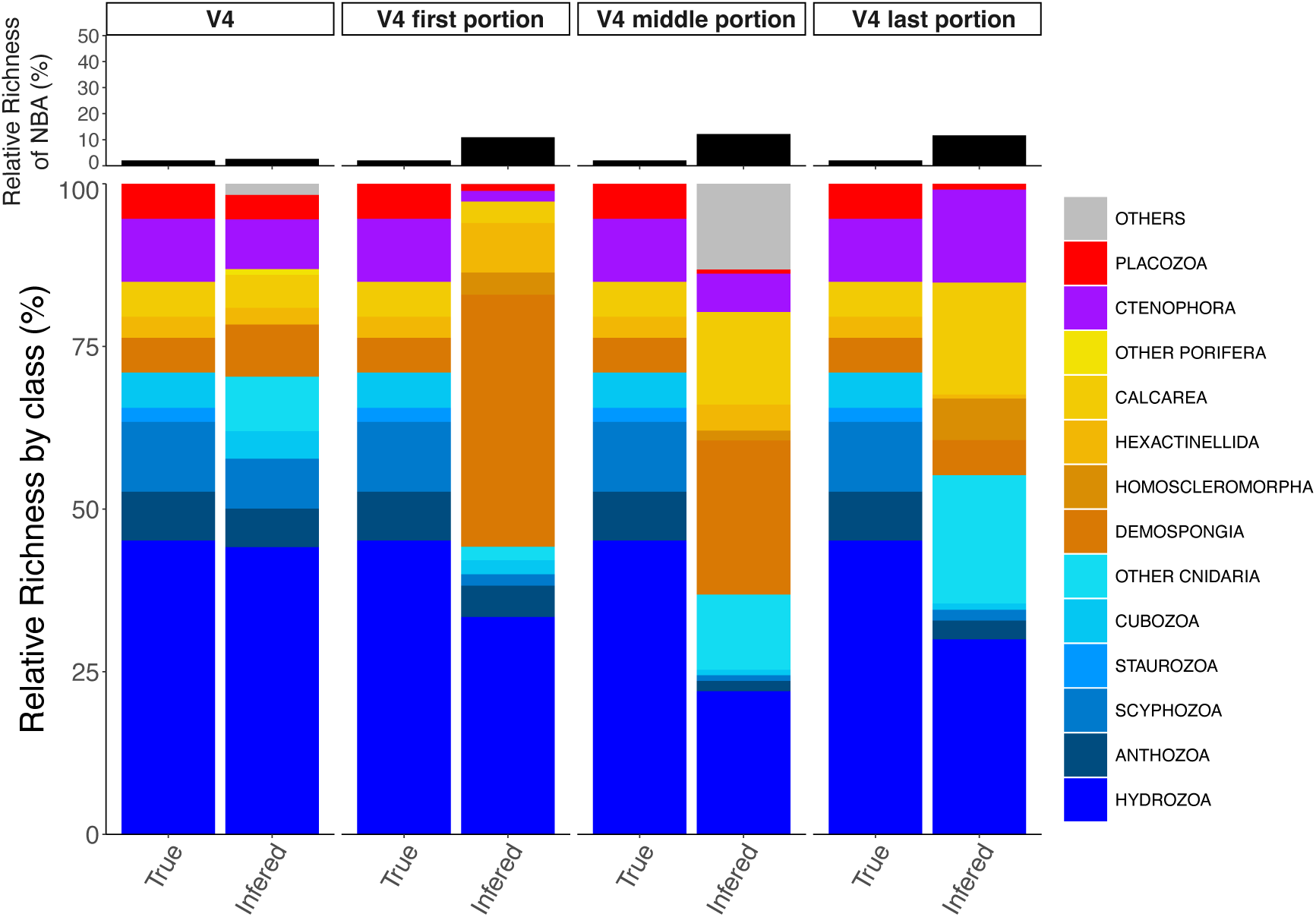
Stacked bar chart exposing the taxonomic analysis of our mock community using the whole V4 amplicon (V4), and three different portions of V4 with the same average length as the V9 region (175 nucleotides). The partition of V4 represents the first, the middle and the last 175 nucleotides portions of the V4 fragment, respectively. For each portion, the true taxonomic identity of the community is compared to the inferred identity using phylogenetic placement and the given barcode. Both the percentage of OTUs assigned as non-bilaterian animals (NBA) over the total number of OTUs (top), and the relative proportion of each non-bilaterian class/phyla (bottom) are shown.

**Supplementary Table 1:** Number of tips by class/phyla in our assembled reference alignments and BTs. For non-targeted taxa (non-animals and Bilateria) broad taxonomic identity of the included sequences is given.

| Class/Phyla | Full BT | V9 BT | V4 BT | Common BT |
| --- | --- | --- | --- | --- |
| Demospongiae | 1037 | 71 | 859 | 55 |
| Hexactinellida | 85 | 7 | 74 | 7 |
| Calcarea | 125 | 17 | 77 | 17 |
| Homoscleromorpha | 24 | 2 | 7 | 2 |
| Ctenophora | 195 | 82 | 108 | 75 |
| Placozoa | 14 | 7 | 12 | 6 |
| Anthozoa | 1118 | 349 | 711 | 228 |
| Hydrozoa | 1447 | 325 | 1011 | 260 |
| Scyphozoa | 228 | 23 | 198 | 22 |
| Staurozoa | 40 | 3 | 35 | 2 |
| Cubozoa | 43 | 19 | 38 | 19 |
| Non-Animals (outgroup) | 20 (1 Plantae, 1 Haptophyta, 1 Rhizaria, 1 Alveolata, 1 Stramenopile, 1 Excavata, 1 Amoebozoa, 1 Nucleariidae, 1 Chytridiomycota, 1 Ascomycota, 1 Blastocladiomycota, 1 Choanozoa, 3 Filasterea, 2 Ichthyosporea, 3 Choanoflagellata) | 8 (1 Plantae, 1 Stramenopile, 1 Haptophyta, 1 Choanoflagellata, 1 Filasterea, 1 Ascomycota, 1 Blastocladiomycota, 1 Choanozoa) | 16 (1 Plantae, 1 Rhizaria, 1 Alveolata, 1 Stramenopile, 1 Excavata, 1 Amoebozoa, 1 Ascomycota, 1 Blastocladiomycota, 1 Choanozoa, 3 Filasterea, 1 Nucleariidae, 2 Ichthyosporea, 1 Choanoflagellata) | 6 (1 Plantae, 1 Filasterea, 1 Ascomycota, 1 Blastocladiomycota, 1 Stramenopile, 1 Choanozoa) |
| Bilateria | 32 (7 Xenacoelomorph | 15 (2 Echinodermata, 4 | 20 (2 Xenacoelomorph | 11 (1 Echinodermata, |
|  | a, 1 Nematoda,<br>1 Hemichordata,<br>2<br>Echinodermata,<br>5 Chordata, 1<br>Nematomorpha,<br>5 Arthropoda, 3<br>Platyhelminthes,<br>2 Mollusca, 4<br>Annelida) | Chordata, 2<br>Platyhelminthes,<br>1 Nematomorpha,<br>2 Arthropoda, 2<br>Annelida, 2<br>Mollusca) | a, 1 Nematoda, 1<br>Echinodermata, 5<br>Chordata, 1<br>Nematomorpha, 4<br>Arthropoda, 1<br>Platyhelminth, 2<br>Mollusca, 3<br>Annelida) | 4 Chordata, 1<br>Platyhelminth, 1<br>Mollusca, 1<br>Annelida, 1<br>Nematomorpha, 2<br>Arthropoda) |
| Total | 4408 | 883 | 3166 | 710 |

**Supplementary Table 2:**
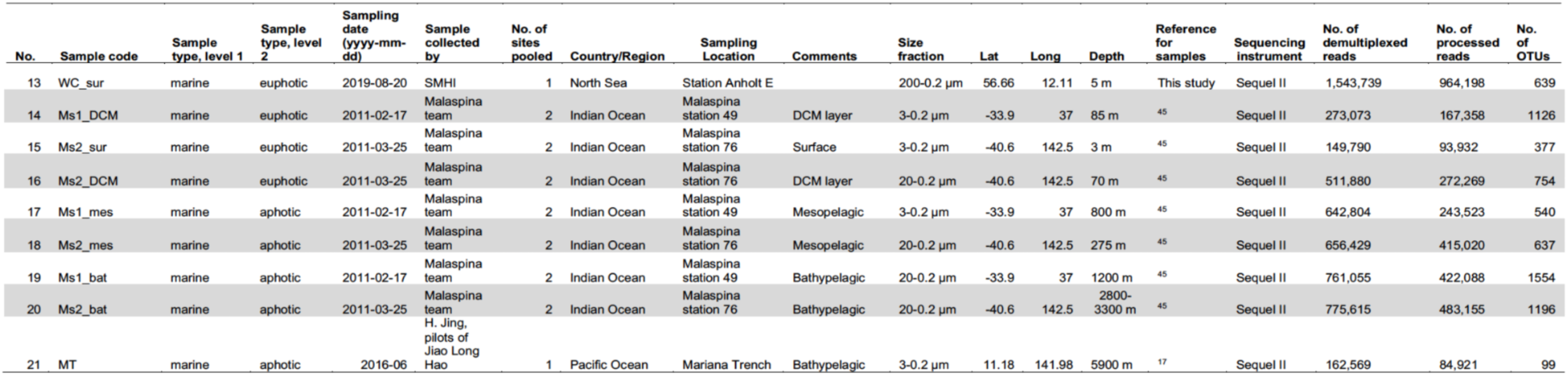
Collection details of the marine samples from Jamy et al., 2022 long-sequence metabarcoding dataset reanalysed here.

All supplementary files can be found in the following Zenodo repository: https://doi.org/10.5281/zenodo.17871426

